# What is a rhythm for the brain? The impact of contextual temporal variability on auditory perception

**DOI:** 10.1101/2023.04.13.536746

**Authors:** Pierre A Bonnet, Mathilde Bonnefond, Anne Kösem

## Abstract

Temporal predictions can be formed and impact perception when sensory timing is fully predictable: for instance, the detection of a target sound is enhanced if it is presented on the beat of an isochronous rhythm. However, natural sensory stimuli, like speech or music, are not entirely predictable, but still possess statistical temporal regularities. We investigated whether temporal expectations can be formed in non-fully predictable contexts, and how the temporal variability of sensory contexts affects auditory perception. Specifically, we asked how “rhythmic” an auditory stimulation needs to be in order to observe temporal predictions effects on auditory discrimination performances. In this behavioral auditory oddball experiment, participants listened to auditory sound sequences where the temporal interval between each sound was drawn from gaussian distributions with distinct standard deviations. Participants were asked to discriminate sounds with a deviant pitch in the sequences. Auditory discrimination performances, as measured with deviant sound discrimination accuracy and response times, progressively declined as the temporal variability of the sound sequence increased. Temporal predictability effects ceased to be observed only for the more variable contexts. Moreover, both global and local temporal statistics impacted auditory perception, suggesting that temporal statistics are promptly integrated to optimize perception. Altogether, these results suggests that temporal predictions can be set up quickly based on the temporal statistics of past sensory events and are robust to a certain amount of temporal variability. Therefore, temporal predictions can be built on sensory stimulations that are not purely periodic nor temporally deterministic.

**Significance statement:** The perception of sensory events is known to be enhanced when their timing is fully predictable. However, it is unclear whether temporal predictions are robust to temporal variability, which is naturally present in many auditory signals such as speech and music. In this behavioral experiment, participants listened to auditory sound sequences where the timing between each sound was drawn from distinct gaussian distributions. Participant’s ability to discriminate deviant sounds in the sequences was function of the temporal statistics of past events: auditory deviant discrimination progressively declined as the temporal variability of the sound sequence increased. Results therefore suggest that auditory perception is sensitive to prediction mechanisms that are involved even if temporal information is not totally predictable.

## Introduction

Temporal predictions are believed to play a key role in the way we process sensory information (Jones, 1976; Schroeder & Lakatos, 2009; Arnal & Giraud, 2012; Nobre et al., 2012). Predicting the timing of future sensory events allows to allocate cognitive resources at the expected time of occurrence, and therefore facilitates the sensory processing of these upcoming stimuli (Large & Jones, 1999; Nobre et al., 1999). As a consequence, the perception of sensory events is improved when their timing is fully predictable. For instance, auditory discrimination is enhanced when the timing of occurrence of the target stimulus is cued (Miniussi et al., 1999; Herbst & Obleser, 2017; Wilsch et al., 2020). Auditory discrimination performances are also improved when the temporal context of the stimulation is deterministic, as when auditory stimulation is periodic (Cravo et al., 2013; Jaramillo & Zador, 2010; Lawrance et al., 2014; Morillon et al., 2016; Rimmele et al., 2011), when temporal intervals are repeated (Breska & Deouell, 2017), or when the temporal intervals are slowing decreasing or increasing at a predictable pace (Cope et al., 2012; Morillon et al., 2016).

However, from a naturalistic point of view, temporal contexts are rarely fully isochronous nor deterministic. Speech acoustic signals in particular presents complex statistical temporal regularities (Singh et al., 2003; Cummins, 2012; Varnet et al., 2017) that are supposedly used to form temporal expectations and influence language comprehension (Tillmann, 2012; Jadoul et al., 2016; Kösem & Van Wassenhove, 2017; Kösem et al., 2018; Aubanel & Schwartz, 2020). How temporal predictions occur in non-fully predictable temporal contexts such as speech and music and how they influence auditory perception is still under debate (Jadoul et al., 2016; Herbst & Obleser, 2017). The aim of this study is to investigate how the temporal statistics of auditory stimulation influences ongoing auditory perception, specifically when the temporal context is not fully predictable. Additionally, the perception of explicit and implicit rhythmicity in audition is known to vary across participants (Geiser et al., 2009; Krause et al., 2010; Repp, 2010) and may have an influence on the way temporal predictions are formed (Doelling & Poeppel, 2015). We therefore also explored whether auditory perception is influenced by the subjective perception of the contextual temporal structure.

To do this, we used an auditory oddball paradigm adapted from the study of Morillon and colleagues (2016). Participants were asked to detect deviant sounds that where embedded in 3 min-long sound sequences. The inter-stimulus-intervals (ISIs) between each sound of the sequence were drawn from Gaussian distributions. The distributions had the same mean (500 ms) but different standard deviations (STD): from 0 ms (periodic) to 150 ms STD (Fig. 1B). Results suggest that (i) temporal predictions can be formed in aperiodic probabilistic context, though auditory detection performance progressively declines with the temporal variability of a context, (ii) these temporal prediction effects are set up quickly from the local temporal statistics of the context. Therefore, this work suggests that temporal prediction mechanisms are robust to temporal variability, and that temporal predictions can be built on sensory stimulations that are not purely periodic nor temporally deterministic.

**Figure 1:**
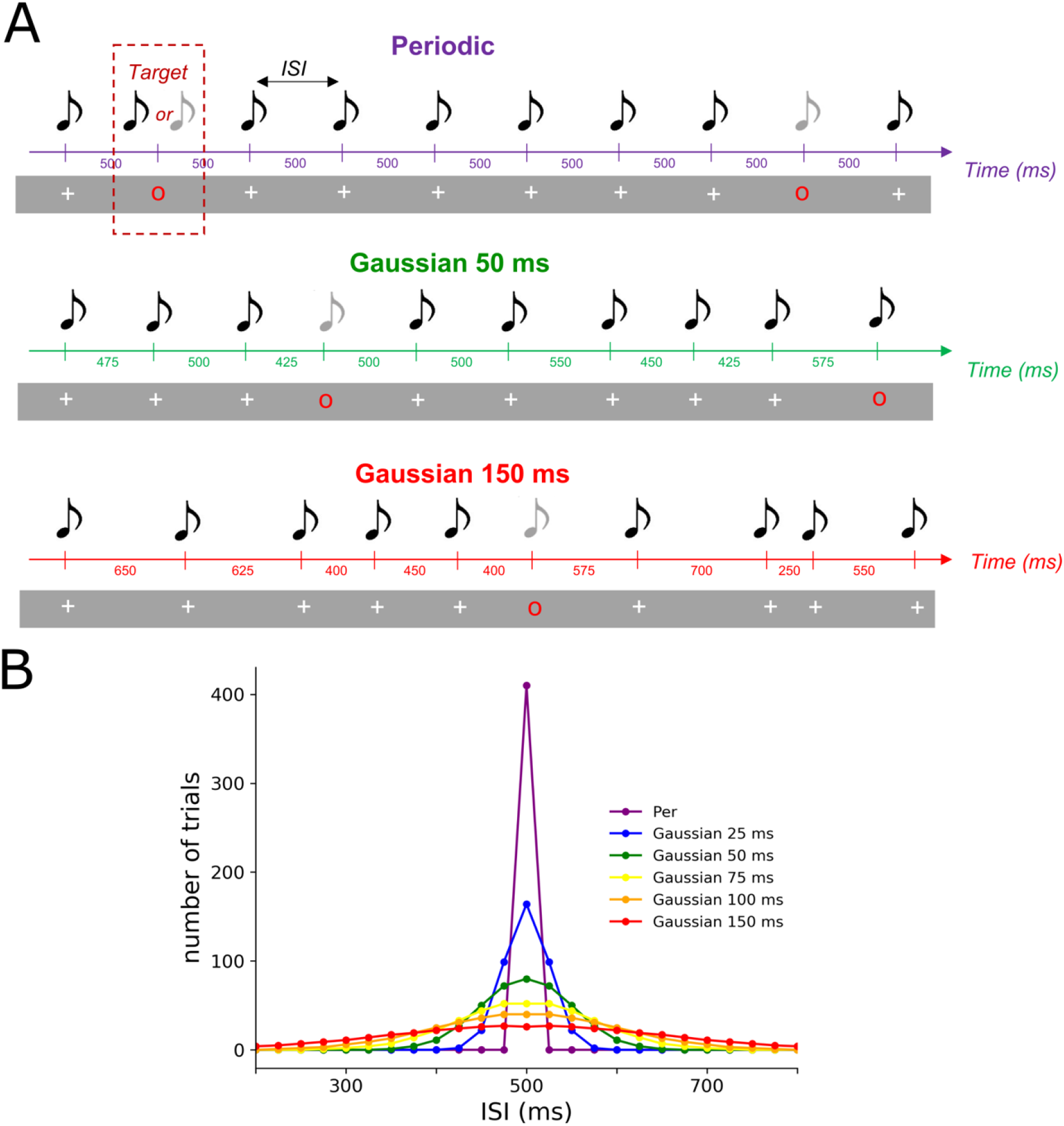
Experimental design. (A) Example of three sequences of different temporal STD used in this experiment. Each sequence consisted of a stream of simultaneous auditory and visual stimuli. A standard stimulus corresponded to a 440 Hz pure tone co-occurring with a white cross. Occasionally a red circle appears in the stream indicating a target stimulus, on which participants had to discriminate between a standard (440 Hz) and deviant (220 Hz) pure tone. (**B**) **Distribution of ISIs in each sequence**. For each sequence, the distributions of the ISIs were drawn of Gaussian distributions with equals means (500 ms) but distinct STDs. Six conditions were designed: from 0 (periodic) to 150 ms of STD with data points built from 100 ms to 900 ms and spaced from 25 ms.

## Materials and Methods

### Participants

Twenty-three participants (11 females, mean age = 25.6 years, 3 left-handed) took part in the experiment. Participants reported no history of neurological or psychiatric disease, normal hearing and normal or corrected-to-normal vision. Four participants had outlier data and were excluded from data analysis: one participant responded at chance level throughout the experiment, three participants had outlier subjective rhythmicity ratings of the sequences (±1.5 interquartile range of regression scores). Therefore, nineteen participants were included for the analysis. The study was approved by an ethical committee (CPP) and all participants signed a written consent and received payment for their participation.

### Stimuli

Participants heard sequences of pure tones, that were either a standard sound (corresponding to a pure 440 Hz sound) or a deviant sound (pure 220 Hz sound). The sounds were presented via headphones for 100 ms (with 5 ms ramp-up and ramp-down in volume). With each sound, visual cues were presented via a digital display (1600, 1024 resolution; 120 Hz refresh rate) and were displayed in front of them (70 cm) in the center of the screen for a duration of 100 ms. Visual cues were synchronized to appear simultaneously with the sounds. A “red circle” visual cue indicated a target and that the participant had to respond to this trial by pressing a button (Fig. 1A). A “white cross” cue indicated a standard trial of the context and that participants did not have to respond to this trial. When the “white cross” was presented on the screen, the synchronized sound was always a standard sound whereas when the “red circle” was presented, the synchronized sound could either be a standard or a deviant sound. In addition to the pure tones, broadband white noise was presented continuously to make the task more difficult. The signal-to-noise ratio between pure tones and white noise was adjusted individually via a staircase procedure (see Procedure). All stimuli were generated and presented via the Psychophysics-3 toolbox.

### Procedure

The experiment was composed of 12 blocks of 3 min 30 s. Each block consisted of a sequence of auditory sensory stimuli masked in constant noise. Participants had to discriminate the sound when a red visual cue was presented on the screen (target trial) (Fig. 1A). To vary the temporal regularities of the context, the Inter-Stimulus-Intervals between the sounds of each block was drawn from distinct distributions. In the Periodic condition, the ISI was fixed at 500 ms. For the Gaussian conditions, the ISIs were drawn from Gaussian distributions with distinct STDs of 25 ms, 50 ms, 75 ms, 100 ms and 150 ms (Fig. 1B). The ISIs data points used to build these gaussian distributions were spaced out every 25 ms to allow accurate sampling of these conditions. Both standards and target trials were drawn from these distributions. Each block consisted of 410 trials including 56 target trials. Between two target trials, a minimum of 4 sounds and a maximum of 10 standard sounds (uniform distribution) could be displayed. The target trials could not appear in the first 10 trials of the sequence. Two blocks were presented for each condition and the block order was pseudo-randomized so that the same condition was not presented twice in succession. Therefore, 112 target trials were obtained for each condition.

After each block, the subjective perception of the rhythmicity of the temporal context was assessed: participants were asked to rate the global rhythmicity of the sequence. We specifically asked to rate whether the sounds in the sequence were presented at a regular pace on a scale from 0 (totally not rhythmic) to 10 (totally periodic). Before the main experiment, a staircase procedure was performed to adjust the broadband noise-to-signal ratio so that the average sound discrimination performance was within ∼80% correct responses. In the staircase, 75 sounds were displayed with periodic ISIs (500 ms) and targets trials could appear every ∼2-3 tones.

### Data analysis

Generalized linear mixed models were computed using lme4 (version 1.1-28) (Bates et al., 2014) with R 4.1.2 (2021-11-01). The dependent variables were the accuracy (binomial distribution) and the subject’s response times (gamma distribution). Temporal STD (continuous variable from 0 ms to 150 ms) and Rhythmicity Rating (continuous variable from 0 to 10) were considered as fixed effects and Subject as a random effect. For accuracy, only the Subject random intercept was added in the model (as adding the random slope did not significantly improve the model’s explained variance). For the response times, both random intercept and random slope were added to the model. Models comparison was done using the likelihood ratio test, and Type II Wald chi-square tests were used to assess significance of fixed effects (Bates et al., 2014; Luke, 2017). We then performed post-hoc tests using the emmeans package version 1.7.4.1, for this we considered Temporal STD as a categorical factor and compared each Temporal STD level (0, 25, 50, 75, 100, 150 ms) using Tukey multiple comparison correction.

We also investigated how the recent temporal statistics in the non-periodic sequences impacted performance (i.e. based on the statistical ISI distribution of the N previous sounds). To do this, we computed for each participant 2-D histograms representing the discrimination accuracy and response times according to the STD and mean ISI of the N previous sounds. Specifically, we computed the mean ISI and STD of the local ISI distribution drawn from the N previous sounds before each target trial (with N ranging from 2 to 7 sounds before target trial). We then binned target trials per ISI distribution mean (from 400 to 600 ms ISI mean, with a sliding window of ±20 ms length) and per ISI distribution STD (from 10 ms to 100 ISI STD, with a sliding window of ±10 ms length). Bins containing less than 5 trials per participant were excluded from further analysis. For each bin, we computed the average accuracy and response time across trials. We obtained a 2-histogram representing how the mean ISI and STD of the 2/3/4/… last sounds impacted accuracy and response times for each participant. We investigated whether, across participants, performances would be relatively better or worse depending on the mean ISI or STD of the local statistics. To test this, we therefore Z-scored the 2-D histograms for each participant and applied cluster-based permutation statistics (using MNE version 1.0.3) to the z-scored data (Maris & Oostenveld, 2007). One sample t-tests against zero were computed for each sample. Adjacent samples with a p-value associated to the t-test of 5% or lower were selected as cluster candidates. The sum of the t-values within a cluster was used as the cluster-level statistic. The reference distribution for cluster-level statistics was computed by performing 1000 random sign-flipping permutations of the data. Clusters were considered significant if the probability of observing a cluster test statistic was below the 2.5-th quantile and above the 97.5-th quantiles for the reference distribution.

## Results

### Temporal variability impacts auditory discrimination accuracy and response times

We tested the effect of the temporal variability of auditory sequences on auditory accuracy and on response times. Participants auditory discrimination significantly decreased as a function of contextual temporal variability (main effect of the factor Temporal STD (*χ2*(1) = 14.574, *p* = 0.0001348)). Discrimination accuracy was highest in the periodic context, and progressively decreased with increasing temporal STD (accuracy decreased by 0.6% every 25 ms). Contrasting each temporal STD condition between one another, post-hoc tests revealed that accuracy in the periodic condition was statistically different from the more aperiodic condition (difference % correct responses Periodic - Gaussian 150 = 4.28%, *p* =0.0061). Moreover, performance was also statistically different between the contexts Gaussian 25 and Gaussian 150 (difference % correct responses Gaussian 25 - Gaussian 150 = 3.76%, *p* =0.0262).). These results suggest that the percentage of correct responses is higher in conditions with less variable contexts even if they are not completely periodic (e.g., in the Gaussian 25 condition) (Fig. 2A).

**Figure 2:**
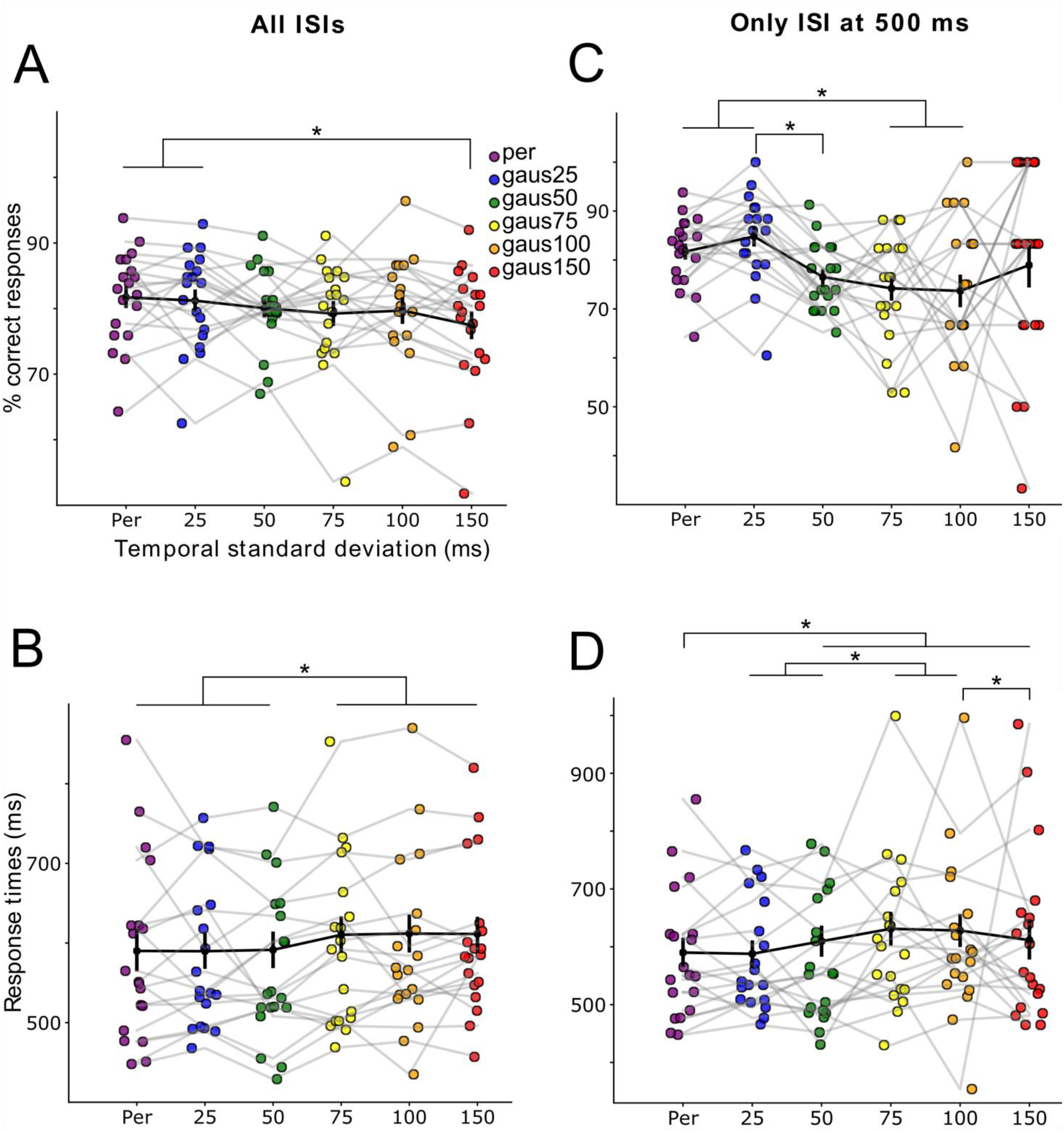
Auditory deviant discrimination is influenced by the temporal variability of the sound sequences. (**A**) Percentage of correct responses and (**C**) Response times as function of the standard deviation of ISIs in the auditory sequences. Each color dot represents a participant. Black dots represent the average across participants. Error bars indicate the Standard Error of the Mean (SEM) and stars indicate significant differences (*p* <0.05). (**B**) Percentage of correct responses and (**D**) Response times restricted to target trials presented at ISI = 500 ms (mean of the distributions).

Response times were also significantly affected by the temporal STD of the context (main effect of the factor Temporal STD (*χ2*(1) = 115.47, *p* < 0.0001)). Response times were faster in temporal contexts with low variability and progressively slowed as the temporal STD increased (response times increased by 5 ms as the context variability increased by 25 ms). Post-hoc tests showed that the three conditions with the lowest temporal variability: Periodic, Gaussian 25 and Gaussian 50 were statistically different from the three conditions with the highest temporal variability: Gaussian 75, Gaussian 100 and Gaussian 150 (difference response times (Periodic, Gaussian 25, Gaussian 50) – (Gaussian 75, Gaussian 100, Gaussian 150) = [-19.3; 22.2 ms], *p* < 0.0001). Results on response times suggest that there is a gap between contexts with low temporal variability and contexts with temporal STD that exceed 75 ms (Fig. 2C).

The more variable the auditory sequence, the more variable the target stimuli’s ISI. It could therefore be possible that the preceding results only reflect the impact of target’s ISI variability, and not of the overall temporal context. In particular, perception is subject to temporal hazard rate, so that auditory discrimination performances improve the longer you wait for the stimulus (Herbst & Obleser, 2019), and reversely, auditory perception performance could decrease drastically for shorter ISIs. To alleviate these effects, we restricted our analyses to all targets whose ISI were of 500 ms only. We still observed similar effects of temporal context as when all ISIs were included. Participants auditory discrimination significantly decreased as a function of contextual temporal variability (main effect of the factor Temporal STD (*χ2*(1) = 14.490, *p* = 0.0001409)). Contrasting each temporal STD condition between one another, post-hoc tests revealed that accuracy in the periodic condition was statistically different from the condition Gaussian 75 ms and from the condition Gaussian 100 ms (difference % correct responses Periodic - Gaussian 75 = 7.50 %, *p* =0.0174; Periodic - Gaussian 100 = 8.04 %, *p* =0.0371). Condition Gaussian 25 ms was also significantly different from the conditions Gaussian 50 ms, Gaussian 75 ms, and Gaussian 100 ms (difference % correct responses Gaussian 25 - Gaussian 50 = 8.27 %; *p* =0.0036; Gaussian 25 - Gaussian 75 = 10.59 %; *p* =0.0004; Gaussian 25 - Gaussian 100 = 11.13 %; *p* =0.0013). These results suggest that when we took only the ISI at the mean of the distributions (500 ms) auditory accuracy was also better in low variability conditions (e.g., periodic and Gaussian 25 ms) compared to conditions with more variability in the context (e.g., Gaussian 75 and 100 ms) (Fig. 2B).

For the response times, there was also a significant main effect of the factor Temporal STD (*χ2*(1) = 77.926; *p* = 2.2e-16). Response times were also faster in temporal contexts with low variability and progressively slowed as the temporal STD increased. Post-hoc tests show that the Periodic condition was different from the Gaussian distributions above 50 ms STD (Periodic – Gaussian 50 = -19.6 ms, *p* < 0.0218; Periodic – Gaussian 75 = -41.1 ms, *p* < 0.0001; Periodic – Gaussian 100 = -38 ms, *p* < 0.0001; Periodic – Gaussian 150 = -21.5 ms, *p* < 0.0218). RTs in Gaussian 25 ms and Gaussian 50 ms were also significantly different from the Gaussian 75 ms and Gaussian 100 ms conditions (Gaussian 25 – Gaussian 75 = -43.3 ms, *p* < 0.0001; Gaussian 25 – Gaussian 100 = -40.2 ms, *p* < 0.0001; Gaussian 50 – Gaussian 75 = -21.4 ms, *p* < 0.0432; Gaussian 50 – Gaussian 100 = -18.3 ms, *p* < 0.0001). RTs in the conditions Gaussian 100 and Gaussian 150 were also statistically different (Gaussian 100 – Gaussian 150 = -16.4 ms, *p* < 0.0288). Results on response times with the ISI at 500 ms only also shows differences between low variability contexts (e.g., Periodic, Gaussian 25 ms or 50 ms) and contexts with more variability in the temporal STD (Fig. 2D).

### Statistical temporal predictions occur rapidly

We further investigated the effect of temporal statistics’ recent history in auditory discrimination performances. Specifically, we computed, across all targets in non-periodic sound sequences, the mean and the STD of the ISI distribution of the N-previous sounds before a target trial (2 to 7 last sounds), and we asked how the temporal statistics of the N previous ISIs impacted the perception of the target trial. When data in all non-periodic sound sequences were aggregated, auditory discrimination performance was sensitive to the mean ISI of the experimental design: discrimination accuracy was highest when the mean ISI of the last N sounds (N being higher or equal to 4) was around 500 ms and was lower when the mean ISI of the last N sounds (N being higher or equal to 4) was below 450 ms and also relatively lower (albeit not significative) above 550 ms (Fig. 3A). Moreover, accuracy was lower when the ISI STD of the last 2-3 sounds was too high (above 50 ms of STD) (Fig. 3D). Similarly, response times were relatively shorter when the mean ISI of the last 2-7 sounds was around 500 ms, thought the effect was not significant (Fig. 3G). Furthermore, when the ISI STD of the last 2-7 sounds was low (around 20-30 ms of STD) responses times were faster, and when the ISI STD was wider (70-90 ms of STD) response times were slower (Fig. 3J).

**Figure 3:**
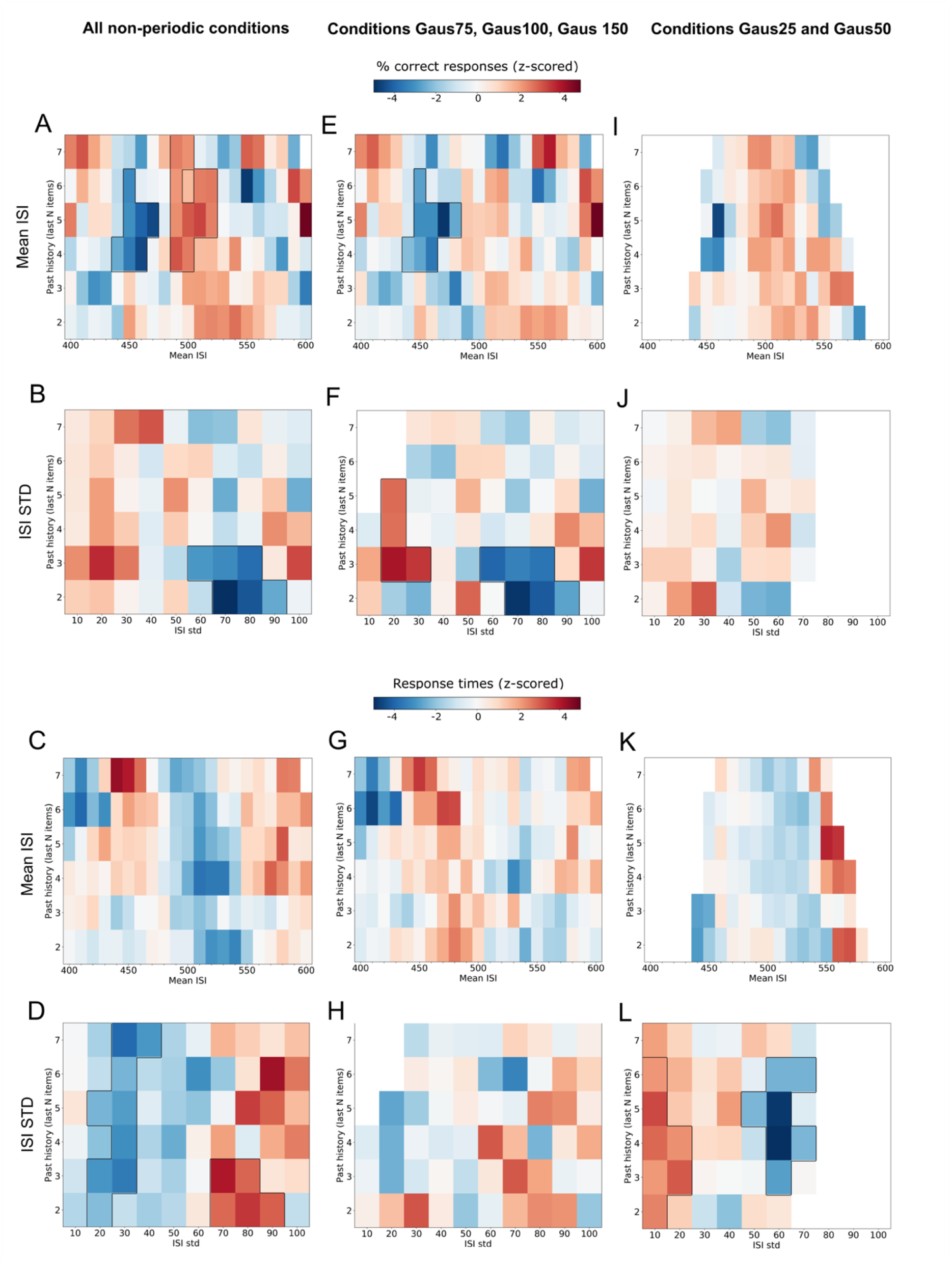
Effect of local temporal ISIs’ statistics on perception. The figures illustrate whether the relative performance of participants was affected by the mean ISI and STD of the previous N sounds. Specifically, we computed the mean ISI and STD of the local ISI distribution drawn from the N previous sounds before each target trial (with N ranging from 2 to 7 sounds before target trial). We then binned target trials per ISI distribution mean (from 400 to 600 ms ISI mean, with a sliding window of ±20 ms length) and per ISI distribution STD (from 10 ms to 100 ISI STD, with a sliding window of ±10 ms length). Bins containing less than 5 trials per participant were excluded from further analysis. For each bin, the average accuracy and response time across trials was computed, and then z-scored across participants. The obtained 2-histograms represent whether accuracy and response times were relatively higher or lower depending on the mean ISI and STD of the 2/3/4/… last sounds. Data were aggregated for either (**A, D, G, J**) all non-periodic contexts, (**B, E, H, K**) the more variable temporal sequences (Gaus75 and higher), and (**C, F, I, L**) the less variable temporal sequences (Gaus25 and Gaus50 conditions). The color label represents the z-scored percentage correct responses or response times. Black lines denote significant clusters. Due to the low variability in conditions Gaus25 and Gaus50 some bins are left white (empty) because the number of trials was not sufficient to be representative (<5).

Data was pooled across all non-periodic sound sequences to maximize the number of trials per bin. We furthered explored the impact of the different global temporal contexts by pooling data from the Gaus25 and Gaus50 conditions, and from the Gaus75, Gaus100 and Gaus150 conditions separately. Pooling the data across the more temporally variable conditions (Gaus75, Gaus100 and Gaus150 conditions), performances followed the same observed patterns as across all non-periodic sound sequences: accuracy was lower when the ISI mean of the previous 4-6 sounds was lower than 450 ms and relatively higher around the mean (although not significant) (Fig. 3B). Moreover, when the ISI STD of the previous sounds was low (20-30 ms) accuracy was higher than for larger STDs (60-90 ms) (Fig. 3E). No significant clusters were found on responses times (Fig. 3HK).

For the less variable conditions (Gaus25 and Gaus50 conditions), we still observed a relatively higher percentage of correct responses, and faster response times around the ISI mean of the distributions (500 ms, albeit no significative cluster was observed) (Fig. 3CI). We observed no conclusive pattern of local ISI STD on accuracy (Fig. 3F). However, we observed different patterns of response times than observed for more temporally variable sequences. Response times were relatively slower when local temporal variability was low (below 20 ms) and faster when the local temporal variability was high (above 60 ms) (Fig. 3L). Please note that the analysis compares the relative change in performance per participant as function of local ISI mean and STD. It is possible that the relative change in performance is less important in low-variable conditions than for the more variable conditions.

Altogether, these results suggest that participants integrated the temporal statistics of the global sound sequences. Furthermore, it suggests that the temporal predictions effects at hand were not related to hazard rate. We also investigated whether recent temporal variability affected performances. Both accuracy and response times were affected by local temporal variability: accuracy was significantly improved, and response times were faster when the local temporal variability was low; conversely accuracy was lower and response times are significantly slower when the local temporal context was highly variable. These findings suggest that temporal expectations form quickly, within a few numbers of sounds in the sequence.

### Link between subjective perception of rhythm and auditory discrimination performance

We also asked participants to subjectively rate their perception of the rhythmicity of each sound sequence. After each sequence, participants rated from 0 (totally arrhythmic) to 10 (totally periodic) the rhythmicity of the sequence of sounds. Participants rated low-variability sequences as more rhythmic: subjective rating was negatively correlated with the temporal variability of context (main effect of the factor Temporal STD (*χ2*(1) = 43.364; *p* < 0.0001)) (Fig. 4A). Temporal STD and Subjective rating being highly correlated, Subjective Rating also correlated with the participant’s discrimination accuracy (main effect of the factor Rating on percentage of correct responses (*χ2*(1) = 14.84; *p* = 0.000117)) (Fig. 4B). Yet, inter-subject variability in rating was observed, with some participants rating non-periodic sequences as more rhythmic than periodic sequences (Fig. 4C). We therefore investigated whether subjective rating could be a predictor of participants performances, specifically whether adding the Subjective Rating as factor with the model fit would explain away more variance in auditory discrimination performance. However, this was not the case: comparing statistical models with the likelihood ratio test, adding the Subjective Rating as a fixed effect in the model did not significantly improve the data fitting for the percentage of correct responses (*p* = 0.072) nor the response times (*p* = 0.624).

**Figure 4:**
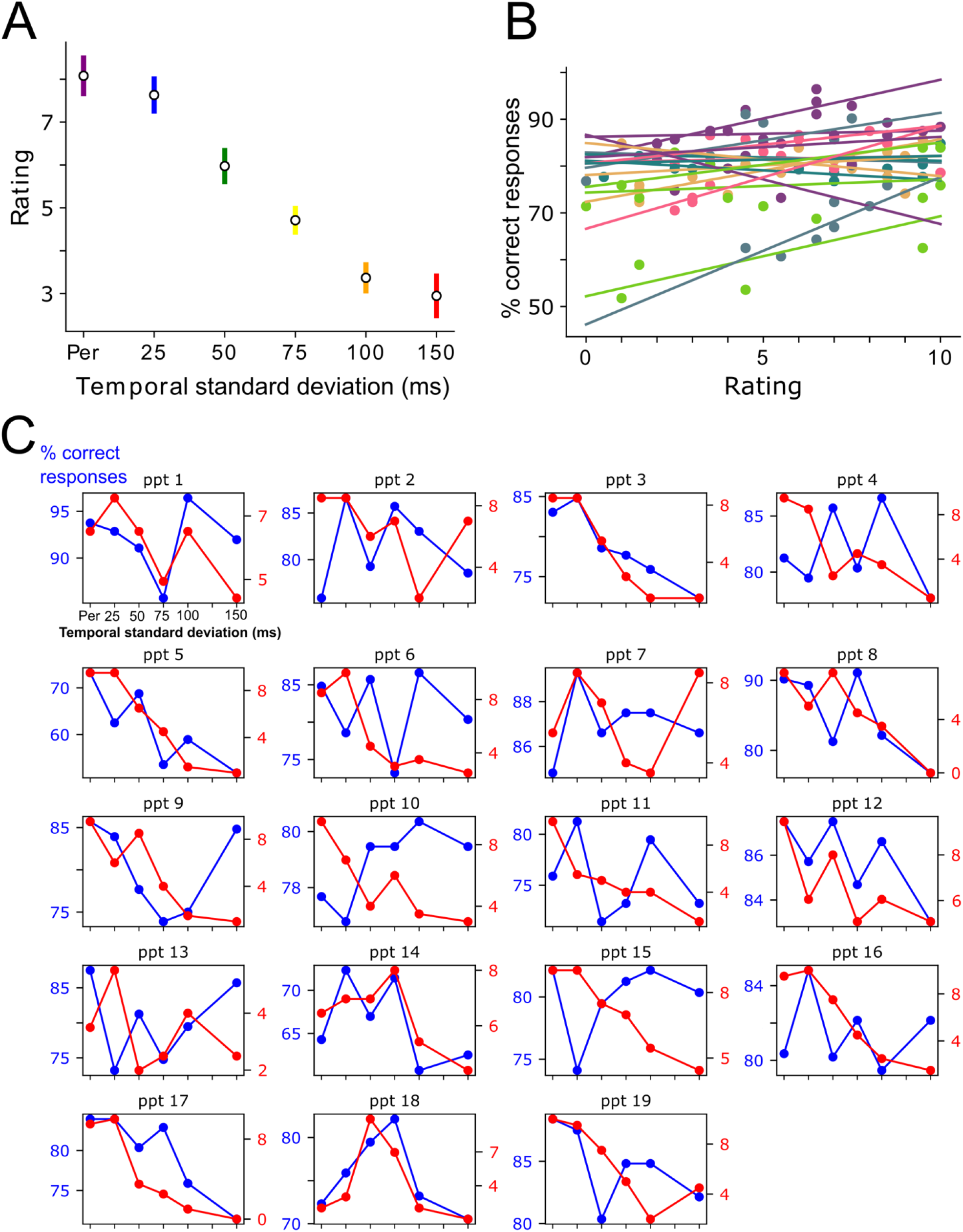
Link between subjective perception of rhythmicity and auditory performances. (**A**) Means of participant’s ratings of the degree of rhythmicity present in the temporal contexts. (**B**) Positive correlation between the rating and the percentage of correct responses. Each point of the same color and the corresponding regression line represent a single participant. (**C**) Individual correlations between percentage of correct responses and rhythmicity ratings. Each figure represents a participant’s data. Blue lines denote the percentage of correct response as a function of temporal standard deviation of sound sequences. Red lines denote the subjective rating of rhythmicity of each sound sequence.

## Discussion

The aim of this study was to investigate the impact of temporal prediction mechanisms on auditory perception in probabilistic temporal contexts. For this, participants were asked to discriminate deviant sounds in auditory sequences, whose ISIs between consecutive sounds were drawn from distinct gaussian distributions. All distributions had the same average ISI (500 ms) but different STDs (from 0 ms up to 150 ms). Auditory perception was influenced by the probabilistic temporal regularities of the sound sequences. Deviant discrimination accuracy was highest and response times were fastest when the deviant sounds were presented in periodic sequences as compared to non-periodic sequences, in line with previous findings (Morillon et al., 2016). However crucially, temporal context also influenced auditory discrimination in the non-periodic sound sequences. Deviant discrimination performances slowly decreased when the temporal variability of the auditory sequences increased. Auditory deviant perception was optimal at average of the ISI distribution of the sequences, suggesting that the timing of target sound was well anticipated based on the probabilistic temporal context. Temporal anticipatory mechanisms were influenced by both global statistics and recent history, as perception was both influenced by the mean ISI of sequences and by the temporal variability of the last sounds prior to target.

### Probabilistic timing influences auditory perception

This study emphasizes that the temporal variability of the context impacts auditory performance. These findings are in line with the literature that shows that auditory perceptual sensitivity is enhanced when stimuli are presented within periodic streams of sensory events (Rimmele et al., 2011; Henry & Obleser, 2012; Cravo et al., 2013; Ten Oever et al., 2014; Morillon et al., 2016; Ten Oever et al., 2017) or when the temporal context is deterministic (Cope et al., 2012; Morillon et al., 2016; Breska & Deouell, 2017). Our findings further suggest that the brain is able to infer events timing in sensory context that have probabilistic temporal properties. Target discrimination accuracy slowly degraded with increasing temporal variability of sound sequences. Response times showed a plateau effect, where a similar increase in response times duration was observed for the more variable contexts from 75 ms STD as compared to contexts with ISI variability below 75 ms STD. When data were restricted to targets presented at the mean ISI of the distribution, we observed that discrimination performances (both in terms of accuracy and RTs) were relatively better for periodic and for low-temporally variable sound sequences as compared to more temporally variable sequences (of standard deviation above 50 ms/75 ms, i.e. 10-15% of the mean ISI of the distribution). Moreover, the persistence of contextual effects when restricting analyses to targets presented at the same ISI (500 ms) further shows that the effect of temporal context on the auditory performances cannot be explained by hazard rate effects alone. Hazard rate implies that performance is solely depends on target’s ISI: performances improve when targets stimuli occur later than expected, and decrease when the sensory stimuli arrive before than expected (Luce, 1986; Nobre & Van Ede, 2018). Here however, we observed temporal contextual effects for the same target ISI.

The effects of probabilistic timing of sensory context have previously been observed on response times (Herbst & Obleser, 2017; Herbst & Obleser, 2019; Grabenhorst et al, 2019; Grabenhorst et al., 2021). In these studies, participants either performed auditory discrimination tasks (Herbst & Obleser, 2017; Herbst & Obleser, 2019), or “set-go” tasks (Grabenhorst et al., 2019; Grabenhorst et al., 2021). The temporal distribution of the foreperiod before the target stimuli influenced response times in both tasks. Importantly, like in this current study, temporal expectancy mechanisms were not uniquely driven by the hazard rate of events, but were also sensitive to the probability of events timing so that response times were fastest when events occur around the mean of contextual temporal probability distribution. Our results further show that not only response times, but also auditory perceptual sensibility is influenced by the temporal probabilities of contextual information. They also highlight that the time required to implement temporal predictions mechanisms based on contextual temporal probabilities is relatively short (temporal statistics from the previous 2-3 ISI can already bias target perception and response times) and depends on the degree of confidence in the temporal regularities of the context.

Knowing that the perception of rhythmic cues in auditory signals is variable across individuals (Potter et al., 2009, Obleser et al., 2017) and could depend on several factors, such as the participant’s musical expertise (Geiser et al., 2009), we examined whether participants accurately perceived the amount of temporal variability in the sound sequences, and whether this impacted their performances. Participants accurately assessed the amount of periodicity in the sound sequences, as their subjective rating of rhythmicity slowly decreased with increased temporal STD of the context. Interestingly, individual variability in the ratings were observed, so that certain participants rated more variable temporal contexts as more rhythmic. However, subjective variations in the perception of rhythmic cues did not significantly add explanatory power to the auditory performances.

### Putative neural mechanisms behind probabilistic temporal predictions

The present findings have implications for current theories and frameworks linking low-frequency neural oscillations to temporal prediction mechanisms in auditory perception. The neural entrainment theory postulates that external rhythms can entrain endogenous neural oscillations, which reflect periodic fluctuations in excitability of neuronal populations (Jones & Boltz, 1989; Large & Jones, 1999; Schroeder & Lakatos, 2009; Cravo et al., 2013). According to this view, neural excitability fluctuations temporally align to the periodic external stream so that the period of high neural excitability coincides with the beat of the external stimuli. A direct prediction of the neural entrainment theory is that perception should be optimal when stimuli are periodic enough to entrain neural oscillations, and when target sensory events occur on beat with the entrained neural oscillation. Here, we do actually report that temporal prediction mechanisms do not only account for purely periodic stimuli but are also robust to a certain amount of temporal variability. Specifically, we found that temporal predictions benefit auditory perception until the variability of temporal context reaches a STD threshold of 10-15% of the mean ISI of the distribution. It is possible that the neural entrainment theory, tested in periodic contexts, could generalize to more complex temporal predictions observed in hierarchically structured rhythms (e.g., speech or music). Importantly, this would explain why temporal properties of speech signals, which are not periodic (Nolan & Jeon, 2014) but are still based on probabilities of occurrence, influence the perceived duration of speech segments and neural dynamics in auditory cortices (Kösem et al., 2018). Interestingly, a recent computational model reports that neural oscillators can handle a certain degree of temporal variability: Stuart–Landau neural oscillatory models are still able to synchronize to temporally variable stimuli, with ISIs drawn from Gaussian distributions with standard deviations going up to 20% of the mean ISI (Doelling & Assaneo, 2021). More research is needed to investigate how observed neural entrainment to auditory stimuli would behave in presence of temporal variability.

Alternatively, the temporal predictions mechanisms observed in periodic and probabilistic contexts could rely on low-frequency dynamics, but would not obviously reflect neural entrainment per se. Evidence for this hypothesis is that low-frequency neural dynamics are shown to reflect temporal predictions in non-entrained sensory context, e.g. when temporal predictions rely on memory-based patterns (Wilsch et al., 2015; Breska & Deouell, 2017; Daume et al., 2021; Herbst et al., 2022). However, it is possible that memory-based predictions and temporal contextual predictions may rely on different co-existing neural mechanisms (Bouwer et al., 2020; Bouwer et al., 2022).

The results of this study also support the view that predictive probabilistic timing is more than hazard rate, and that it also relies on the probability density function of the timing of previous sensory events. As such, the mechanisms related to hazard rate and contextual temporal predictions could be dissociated and have different mechanistic origins. While hazard rate processing seems to rely on motor regions (Herbst & Obleser, 2019; Cui et al., 2009), contextual probabilistic timing may involve a different distributed neural architecture, including early sensory areas (Bueti et al., 2010; Herbst & Obleser, 2019).

Taken together, the results of this study show that temporal predictive mechanisms influence auditory perception in implicit probabilistic temporal contexts, and that they are robust to some amount temporal variability.

## Acknowledgments

This work was supported by a Marie Sklodowska-Curie Fellowship (grant 843088) and by an ANR grant (ANR-21-CE37-0003) to A.K. We would like to thank Oussama Abdoun for his assistance with the data analysis.

## References

Arnal, L. H., & Giraud, A.-L. (2012). Cortical oscillations and sensory predictions. Trends in Cognitive Sciences, 16(7), 390–398. https://doi.org/10.1016/j.tics.2012.05.003

Aubanel, V., & Schwartz, J.-L. (2020). The role of isochrony in speech perception in noise. Scientific Reports, 10(1), 19580. https://doi.org/10.1038/s41598-020-76594-1

Bates, D., Mächler, M., Bolker, B., & Walker, S. (2014). Fitting Linear Mixed-Effects Models using lme4 (1406.5823). arXiv. https://doi.org/10.48550/arXiv.1406.5823

Bouwer, F. L., Honing, H., & Slagter, H. A. (2020). Beat-based and memory-based temporal expectations in rhythm: similar perceptual effects, different underlying mechanisms. Journal of Cognitive Neuroscience, 32(7), 1221–1241.

Bouwer, F. L., Fahrenfort, J. J., Millard, S. K., Kloosterman, N. A., & Slagter, H. A. (2022). A silent disco: Persistent entrainment of low-frequency neural oscillations underlies beat-based, but not pattern-based temporal expectations. bioRxiv, 2020-01.

Breska, A., & Deouell, L. Y. (2017). Neural mechanisms of rhythm-based temporal prediction: Delta phase-locking reflects temporal predictability but not rhythmic entrainment. PLOS Biology, 15(2), e2001665. https://doi.org/10.1371/journal.pbio.2001665

Bueti, D., Bahrami, B., Walsh, V., & Rees, G. (2010). Encoding of temporal probabilities in the human brain. Journal of Neuroscience, 30(12), 4343–4352.

Cope, T. E., Grube, M., & Griffiths, T. D. (2012). Temporal predictions based on a gradual change in tempo. The Journal of the Acoustical Society of America, 131(5), 4013–4022. https://doi.org/10.1121/1.3699266

Cravo, A. M., Rohenkohl, G., Wyart, V., & Nobre, A. C. (2013). Temporal Expectation Enhances Contrast Sensitivity by Phase Entrainment of Low-Frequency Oscillations in Visual Cortex. Journal of Neuroscience, 33(9), 4002–4010. https://doi.org/10.1523/JNEUROSCI.4675-12.2013

Cui, X., Stetson, C., Montague, P. R., & Eagleman, D. M. (2009). Ready… go: amplitude of the fMRI signal encodes expectation of cue arrival time. PLoS biology, 7(8), e1000167.

Cummins, F. (2012). Oscillators and Syllables: A Cautionary Note. Frontiers in Psychology, 3. https://doi.org/10.3389/fpsyg.2012.00364

Daume, J., Wang, P., Maye, A., Zhang, D., & Engel, A. K. (2021). Non-rhythmic temporal prediction involves phase resets of low-frequency delta oscillations. NeuroImage, 224, 117376. https://doi.org/10.1016/j.neuroimage.2020.117376

Doelling, K. B., & Poeppel, D. (2015). Cortical entrainment to music and its modulation by expertise. Proceedings of the National Academy of Sciences, 112(45), E6233–E6242.

Doelling, K. B., & Assaneo, M. F. (2021). Neural oscillations are a start toward understanding brain activity rather than the end. PLoS biology, 19(5), e3001234.

Geiser, E., Ziegler, E., Jancke, L., & Meyer, M. (2009). Early electrophysiological correlates of meter and rhythm processing in music perception. Cortex, 45(1), 93–102. https://doi.org/10.1016/j.cortex.2007.09.010

Giraud, A.-L., & Poeppel, D. (2012). Cortical oscillations and speech processing: Emerging computational principles and operations. Nature Neuroscience, 15(4), 511–517. https://doi.org/10.1038/nn.3063

Grabenhorst, M., Michalareas, G., Maloney, L. T., & Poeppel, D. (2019). The anticipation of events in time. Nature Communications, 10(1), 5802. https://doi.org/10.1038/s41467-019-13849-0

Grabenhorst, M., Maloney, L. T., Poeppel, D., & Michalareas, G. (2021). Two sources of uncertainty independently modulate temporal expectancy. Proceedings of the National Academy of Sciences, 118(16), Article 16. https://doi.org/10.1073/pnas.2019342118

Henry, M. J., & Obleser, J. (2012). Frequency modulation entrains slow neural oscillations and optimizes human listening behavior. Proceedings of the National Academy of Sciences, 109(49), 20095–20100.

Herbst, S. K., & Obleser, J. (2017). Implicit variations of temporal predictability: Shaping the neural oscillatory and behavioural response. Neuropsychologia, 101, 141–152. https://doi.org/10.1016/j.neuropsychologia.2017.05.019

Herbst, S. K., & Obleser, J. (2019). Implicit temporal predictability enhances pitch discrimination sensitivity and biases the phase of delta oscillations in auditory cortex. NeuroImage, 203, 116198.

Herbst, S. K., Stefanics, G., & Obleser, J. (2022). Endogenous modulation of delta phase by expectation–A replication of Stefanics et al., 2010. Cortex, 149, 226–245. https://doi.org/10.1016/j.cortex.2022.02.001

Jadoul, Y., Ravignani, A., Thompson, B., Filippi, P., & de Boer, B. (2016). Seeking Temporal Predictability in Speech: Comparing Statistical Approaches on 18 World Languages. Frontiers in Human Neuroscience, 10. https://www.frontiersin.org/articles/10.3389/fnhum.2016.00586

Jaramillo, S., & Zador, A. (2010). Auditory cortex mediates the perceptual effects of acoustic temporal expectation. Nature Precedings, 1–1. https://doi.org/10.1038/npre.2010.5139.1

Jones, M. R. (1976). Time, our lost dimension: Toward a new theory of perception, attention, and memory. Psychological Review, 83(5), 323–355. https://doi.org/10.1037/0033-295X.83.5.323

Jones, M. R., & Boltz, M. (1989). Dynamic attending and responses to time. Psychological Review, 96(3), 459–491. https://doi.org/10.1037/0033-295X.96.3.459

Kösem, A., & Van Wassenhove, V. (2017). Distinct contributions of low-and high-frequency neural oscillations to speech comprehension. Language, cognition and neuroscience, 32(5), 536–544.

Kösem, A., Bosker, H. R., Takashima, A., Meyer, A., Jensen, O., & Hagoort, P. (2018). Neural entrainment determines the words we hear. Current Biology, 28(18), 2867–2875.

Krause, V., Pollok, B., & Schnitzler, A. (2010). Perception in action: The impact of sensory information on sensorimotor synchronization in musicians and non-musicians. Acta Psychologica, 133(1), 28–37. https://doi.org/10.1016/j.actpsy.2009.08.003

Large, E. W., & Jones, M. R. (1999). The dynamics of attending: How people track time-varying events. Psychological Review, 106(1), 119–159. https://doi.org/10.1037/0033-295X.106.1.119

Lawrance, E. L. A., Harper, N. S., Cooke, J. E., & Schnupp, J. W. H. (2014). Temporal predictability enhances auditory detection. The Journal of the Acoustical Society of America, 135(6), EL357–EL363. https://doi.org/10.1121/1.4879667

Luce, R. D. (1986). Response times: Their role in inferring elementary mental organization (No. 8). Oxford University Press on Demand.

Luke, S. G. (2017). Evaluating significance in linear mixed-effects models in R. Behavior Research Methods, 49(4), 1494–1502. https://doi.org/10.3758/s13428-016-0809-y

Maris, E., & Oostenveld, R. (2007). Nonparametric statistical testing of EEG-and MEG-data. Journal of neuroscience methods, 164(1), 177–190.

Miniussi, C., Wilding, E. L., Coull, J. T., & Nobre, A. C. (1999). Orienting attention in time: Modulation of brain potentials. Brain, 122(8), 1507–1518.

Morillon, B., Schroeder, C. E., Wyart, V., & Arnal, L. H. (2016). Temporal Prediction in lieu of Periodic Stimulation. Journal of Neuroscience, 36(8), 2342–2347. https://doi.org/10.1523/JNEUROSCI.0836-15.2016

Nobre, A. C., Coull, J. T., Frith, C. D., & Mesulam, M. M. (1999). Orbitofrontal cortex is activated during breaches of expectation in tasks of visual attention. Nature Neuroscience, 2(1), 11–12. https://doi.org/10.1038/4513

Nobre, A. C., Rohenkohl, G., & Stokes, M. G. (2012). Nervous anticipation: Top-down biasing across space and time. In Cognitive neuroscience of attention, 2nd ed (pp. 159–186). The Guilford Press.

Nobre, A. C., & Van Ede, F. (2018). Anticipated moments: temporal structure in attention. Nature Reviews Neuroscience, 19(1), 34–48.

Nolan, F., & Jeon, H.-S. (2014). Speech rhythm: A metaphor? Philosophical Transactions of the Royal Society B: Biological Sciences, 369(1658), 20130396. https://doi.org/10.1098/rstb.2013.0396

Obleser, J., Henry, M. J., & Lakatos, P. (2017). What do we talk about when we talk about rhythm? PLOS Biology, 15(9), e2002794. https://doi.org/10.1371/journal.pbio.2002794

Potter, D. D., Fenwick, M., Abecasis, D., & Brochard, R. (2009). Perceiving rhythm where none exists: Event-related potential (ERP) correlates of subjective accenting. Cortex, 45(1), 103–109. https://doi.org/10.1016/j.cortex.2008.01.004

Repp, B. H. (2010). Sensorimotor synchronization and perception of timing: Effects of music training and task experience. Human Movement Science, 29(2), 200–213. https://doi.org/10.1016/j.humov.2009.08.002

Rimmele, J., Jolsvai, H., & Sussman, E. (2011). Auditory Target Detection Is Affected by Implicit Temporal and Spatial Expectations. Journal of Cognitive Neuroscience, 23(5), 1136–1147. https://doi.org/10.1162/jocn.2010.21437

Schroeder, C. E., & Lakatos, P. (2009). Low-frequency neuronal oscillations as instruments of sensory selection. Trends in Neurosciences, 32(1), 9–18. https://doi.org/10.1016/j.tins.2008.09.012

Singh, N. C., & Theunissen, F. E. (2003). Modulation spectra of natural sounds and ethological theories of auditory processing. The Journal of the Acoustical Society of America, 114(6), 3394–3411.

Ten Oever, S., Schroeder, C. E., Poeppel, D., van Atteveldt, N., & Zion-Golumbic, E. (2014). Rhythmicity and crossmodal temporal cues facilitate detection. Neuropsychologia, 63, 43–50. https://doi.org/10.1016/j.neuropsychologia.2014.08.008

Ten Oever, S., Schroeder, C. E., Poeppel, D., Atteveldt, N. van, Mehta, A. D., Mégevand, P., Groppe, D. M., & Zion-Golumbic, E. (2017). Low-Frequency Cortical Oscillations Entrain to Subthreshold Rhythmic Auditory Stimuli. Journal of Neuroscience, 37(19), 4903–4912. https://doi.org/10.1523/JNEUROSCI.3658-16.2017

Tillmann, B. (2012). Music and Language Perception: Expectations, Structural Integration, and Cognitive Sequencing. Topics in Cognitive Science, 4(4), 568–584. https://doi.org/10.1111/j.1756-8765.2012.01209.x

Varnet, L., Ortiz-Barajas, M. C., Erra, R. G., Gervain, J., & Lorenzi, C. (2017). A cross-linguistic study of speech modulation spectra. The Journal of the Acoustical Society of America, 142(4), 1976–1989. https://doi.org/10.1121/1.5006179

Wilsch, A., Henry, M. J., Herrmann, B., Maess, B., & Obleser, J. (2015). Slow-delta phase concentration marks improved temporal expectations based on the passage of time. Psychophysiology, 52(7), 910–918. https://doi.org/10.1111/psyp.12413

Wilsch, A., Mercier, M. R., Obleser, J., Schroeder, C. E., & Haegens, S. (2020). Spatial attention and temporal expectation exert differential effects on visual and auditory discrimination. Journal of Cognitive Neuroscience, 32(8), 1562–1576.

